# How do ecological and social environments reflect parental roles in birds? A comparative analysis

**DOI:** 10.1101/2020.12.24.424295

**Authors:** Xiaoyan Long, Yang Liu, András Liker, Franz J. Weissing, Jan Komdeur, Tamás Székely

## Abstract

Parental roles are highly diverse in animal taxa. Since caring is an important determinant of fitness, understanding the origin and maintenance of various parental care strategies is a key question in evolutionary biology. Here we investigate parental care patterns in birds, which exhibit a remarkable diversity of parental sex roles. By means of phylogenetically informed comparative analyses we investigate whether and how care provisioning is predicted by ecology and social environment. Making use of the most comprehensive dataset including 1101 species that represent 126 avian families, we show that sex differences in parental care are neither related to food type nor to nest type, two key ecological factors. However, we found an effect of the social environment, as males tend to care relatively more in in colonial species than in non-colonial species. Taken together, these results highlight the importance of social effects for evolution of diverse parental sex roles.

## INTRODUCTION

Parental care, which increases offspring survival at the cost of parents’ viability and fecundity, varies widely among animal taxa (Balshine 2012; Trumbo 2012). Birds are characterized by an extraordinary diversity in parental roles, ranging from female-only care (e.g., hummingbirds, where only the females build nests, incubate eggs and feed the young), to biparental care (e.g., woodpeckers, where the two sexes share parental duties), and male-only care (e.g., some shorebirds, where the males incubate the eggs and rear the young without any support by the females) (Cockburn 2006; Remeš et al. 2015). Understanding this diversity is important as parental roles have repercussions on many other characteristics, such as sex differences in morphology, demography and mating strategies (Emlen and Oring 1977; Fairbairn et al. 2007; Royle et al. 2012; Klug 2018).

Various ecological and social variables have been proposed to influence parental sex roles (McGraw et al. 2010; Klug et al. 2012). Firstly, it has been proposed that food type is related to sex differences in parental care. It has been argued that uniparental care is to be expected in bird species feeding on plant materials such as fruits and nectar; as such food sources tend to be seasonally abundant, one parent should suffice to efficiently provision the young (Cockburn 2006; Barve and La Sorte 2016). In contrast, biparental care might be expected in bird species feeding on insects or other animals; such food is often dispersed and difficult to catch and both parents are required to satisfy the demands of their offspring (Badyaev and Ghalambor 1998). However, this line of argumentation is not fully convincing. Since plant food is nutritionally inferior to animal food, herbivorous chicks may need much larger amounts of food to match their energetic demands; accordingly, both parents might be required to collect or defend enough food. Conversely, a single parent might be sufficient to satisfy the demands of carnivorous offspring if only few highly nutritious food items need to be collected per time unit.

Secondly, it has been proposed that nest type (open or closed) is an important determinant of parental sex roles (Collias and Collias 1984; Hansell 2000). Open nests are exposed to the environment and consequently face a high risk of predation (Collias 1964; Cody 1966; Martin 1995; Lima 2009). It has been argued that this will select for biparental care, since the predation risk can be reduced considerably if both parents are around (Montgomerie and Weatherhead 1988; Kleindorfer and Hoi 1997; Martin and Briskie 2009). In contrast, uniparental care might be expected to be common in species with closed nests, which provide a good protection from predators. However, an argument could also be made for the opposite pattern. For example, the presence of both parents (and, in particular, the presence of a brightly coloured father) could make an open nest more conspicuous to predators; hence predation risk might be enhanced rather than reduced if both parents are around (Skutch 1949; Martin et al. 2000). Predation pressure is generally lower in species with closed nests, but such species tend to have greater clutch sizes than those building open nests (Jetz et al. 2008). Accordingly, just in cavity breeders both parents may be needed to meet the energetic demands of the offspring.

Thirdly, social environment is also predicted to affect parental care strategies. In colonial breeding species, pairs live in groups and often in high density, which reduces the risk of predation and allows for exchanging information about food resources (Alexander 1974; Perrins and Birkhead 1983; Krause and Ruxton 2002; Evans et al. 2016). Accordingly, one might expect that uniparental care is more prevalent in colonially breeding species than in solitarily breeding species. In addition, opportunities for extra-pair copulations are typically high in colonial species as breeding density is high, lowering the certainty of paternity (Westneat and Sherman, 1997; Mayer and Pasinelli 2013) and selecting for reduced paternal care. But again, a case could also be made for a different prediction. Colonially breeding species almost invariable face high intraspecific competition (interference competition and competition for food and other resources, Perrins and Birkhead 1983; Krause and Ruxton 2002). Based on this, one would expect biparental care, as both parents are needed to successfully raise the young.

In view of all this, it is not self-evident which parental care patterns are to be expected under different ecological and social scenarios. It is therefore important to investigate what the data say. To what extent and in what way do ecological conditions (like food quality or predation pressure) or the social environment (like coloniality) reflect parental roles? To answer this question, we here apply phylogenetic comparative analyses to a comprehensive dataset including 1101 avian species (26 orders and 119 families). First, we examine whether parental roles are associated with food type. Specifically, we investigate whether plant-eating species exhibit uniparental or sex-biased parental care while carnivorous species exhibit more biparental care. Second, we explore whether parental care is associated with nest type, where nest type is viewed as a proxy for the risk of predation. Specifically, we test whether open nesters or closed nesters are more likely to provide biparental care. Third, we study whether parental care patterns are associated with coloniality. Specifically, we test whether colonial breeders tend to exhibit female-biased care while parental sex roles are less biased in solitarily breeding bird species.

## METHODS

### Data collection

We collected data from reference works (e.g., The Birds of the Western Palearctic, The Birds of North America, Handbook of Australian, New Zealand and Antarctic Birds), preexisting datasets (see below) and primary literatures by using Web of Science and Google Scholar. We added more species with available data on parental behavior to an existing dataset used by Liker et al. (2015). Then we augmented the dataset with expanded information on parental roles by extracting ecological and social traits (food type, nest type and coloniality). The final dataset included 1101 species (26 orders and 119 families) representing a broad spectrum of avian taxa. All data are available in the supplementary materials.

### Parental care variables

Bird species exhibit diverse forms of parental care, ranging from the preparation of the nest to nutrition provision. Here, we investigate eight types of avian parental behavior: nest building, nest guarding, incubation, chick brooding, chick feeding, chick guarding, post-fledgling feeding and post-fledgling guarding. Since quantitative data on parental contribution were not available for many species, we scored each care type on a 5-point scale: −1: no male care; −0.5: 1–33% male care; 0: 34– 66% male care; 0.5: 67–99% male care; 1: 100% male care. When quantitative data were not measured, we used the information from verbal descriptions. For instance, species with more male contribution to nestling feeding would be scored 0.5 on chick feeding. By means of the scoring system, the estimates of paternal contribution and maternal contribution were fully dependent, that is, male scores would always be the additive inverse of female scores. Therefore, the scores directly reflected the sex differences in parental roles. 0 indicated approximately equal parental investment by both parents, 0.5 and −0.5 represented male-biased and female-biased parental contribution, respectively, 1 and −1 suggested male-only care and female-only care, respectively.

We then divided parental activities into two breeding phases: (i) pre-hatching care, which involved nest building, nest guarding and incubation and (ii) post-hatching care, which included chick brooding, chick feeding, chick guarding, post-fledgling feeding and post-fledgling guarding. To score the relative participation in pre-hatching care and post-hatching care by males, we calculated the mean scores of different components of parental behavior for each state. The relative participation in pre-hatching care by males strongly correlated with the relative participation in nest building, incubation and nest guarding by males (*r^2^* = 0.508-0.644, *P* < 0.001, see details in Supplementary Figure S1 and Table S1). Similarly, the relative participation in post-hatching care by males was significantly related to the relative participation in chick brooding, chick feeding, chick guarding, post-fledgling feeding and post-fledgling guarding by males (*r^2^* = 0.393-0.729, *P* < 0.001, see details in Supplementary Figure S1 and Table S1). This suggested that pre- and post-hatching care can reliably represent a set of specific care components.

### Ecological and social variables

The diet of bird species was classified into two categories: (i) plant materials which included fruit, seed, leaves and (ii) animals which included invertebrates and vertebrates. For omnivorous species, their mainly eaten food category was allocated. Since parents and nestlings subsist on different food items in some species (e.g., In willow ptarmigan (*Lagopus lagopus*) the adults forage for plant materials all year round while their nestlings are usually fed on insects (Peters 1958)), we collected diet of parents and nestlings separately.

Nest type was treated as binary variables (open or closed). Open nests, which are exposed to adverse weather conditions and predators, included scrapes (e.g., nests of many shorebirds), cups (e.g., nests of many passerines) and platforms (e.g., nests of raptors) (Hansell 2000). Closed nests are completely covered by the walls or pliable materials, that is, they can only be accessed by the small entrance. For instance, cavities (e.g., nests of woodpeckers), burrows (e.g., nests of many seabirds), domes and globes (e.g., nests of weavers) are all enclosed structures (Hansell 2000). We only extracted data on nest type from studies of natural nests (i.e., nest-box studies were excluded).

Coloniality was categorized into (i) solitary breeding, breeders never live in groups, (ii) semi-colonial breeding, breeders are either solitary or colonial, and (iii) colonial breeding, individuals always breed in groups and they defend a territory which only consists of the nest sites (Perrins and Birkhead, 1983, Van Turnhout et al. 2010). We only extracted data on coloniality from studies of natural nests, since the studies of nest-box artificially changed the spatial distribution of nests.

### Phylogenetic analyses

To test whether pre-hatching care differs from post-hatching care, we conducted phylogenetic paired t-tests (Lindenfors et al. 2010). We first estimated the corresponding phylogenetic mean value of pre-hatching and post-hatching care of each species, then compared whether the mean difference was different from zero (Lindenfors et al. 2010). The analyses were implemented in R (3.4.2) using the ‘phytools’ package (Revell 2012).

We analyzed the correlation between parental care variables and predictor variables by using phylogenetic generalized least squares (PGLS) (Freckleton et al. 2002). This technique controls for the dependence of species traits as a result of shared evolutionary history by estimating the expected covariance structure, then modified slope and intercept estimates would be calculated. In all analyses, Pagel’s λ which varies between 0 and 1 was estimated to represent the phylogenetic signal (Freckleton et al. 2002). A trait with strong phylogenetic signal is more similar among closely related species, while data points are more independent if phylogenetic signal is weak.

Considering the uncertainty of phylogenetic estimation caused by the absence of empirical support on the prediction of evolutionary relationships among species (Jetz et al. 2012), we randomly extracted 100 phylogenetic trees from the most comprehensive avian phylogenies (Jetz et al. 2012). Each PGLS model was analyzed across all of the trees and the mean value of resulting 100 parameter estimates were calculated.

For each dependent variable (i.e., the relative participation in pre-hatching care by males, the relative participation in post-hatching care by males), we established separate PGLS models to investigate the effect of each ecological and social traits. Here, we present (1) the results of bivariate models which only included one of the main predictors, and (2) the results of multi-predictor models. Since nestling diet was related to parental diet, two multi-predictor PGLS models were built in order to avoid the problem of multicollinearity. These two multi-predictor models contained the following predictors i) parental diet, nest type and coloniality; ii) nestling diet, nest type and coloniality. Moreover, two crucial life-history traits were included in all multi-predictor models: body mass (log-transformed) and chick development mode (precocial vs. altricial). This is due to the fact that body mass strongly correlated with several life-history traits (e.g., longevity, Lindstedt and Calder 1976), and chick development was suggested to affect parental roles (Thomas and Székely 2005; Olson et al. 2008). All PGLS analyses were carried out in the R statistical computing environment as well, using the package “caper” (Orme 2013).

## RESULTS

### Pre-hatching care vs. post-hatching care

Birds exhibit diverse parental roles, consistently with expectations, including female-only care, biparental care and male-only care (Figure 1). Even in the same clade, different parental care patterns can be observed (Figure 1a-d), corresponding to the intermediate phylogenetic signal of each breeding activities (λ = 0.187-0.755, see statistical estimates in Supplementary Table S1). For instance, uniparental care by the male or the female, and biparental care coexist in Anseriformes, Charadriiformes, Procellariiformes, Psittaciformes and Passeriformes. A remarkable diversity is observed in shorebirds in that all care types (female-only care, female-biased care, biparental care, male-biased care and male-only care) can be found (Figure 1c, d).

**Figure 1.**
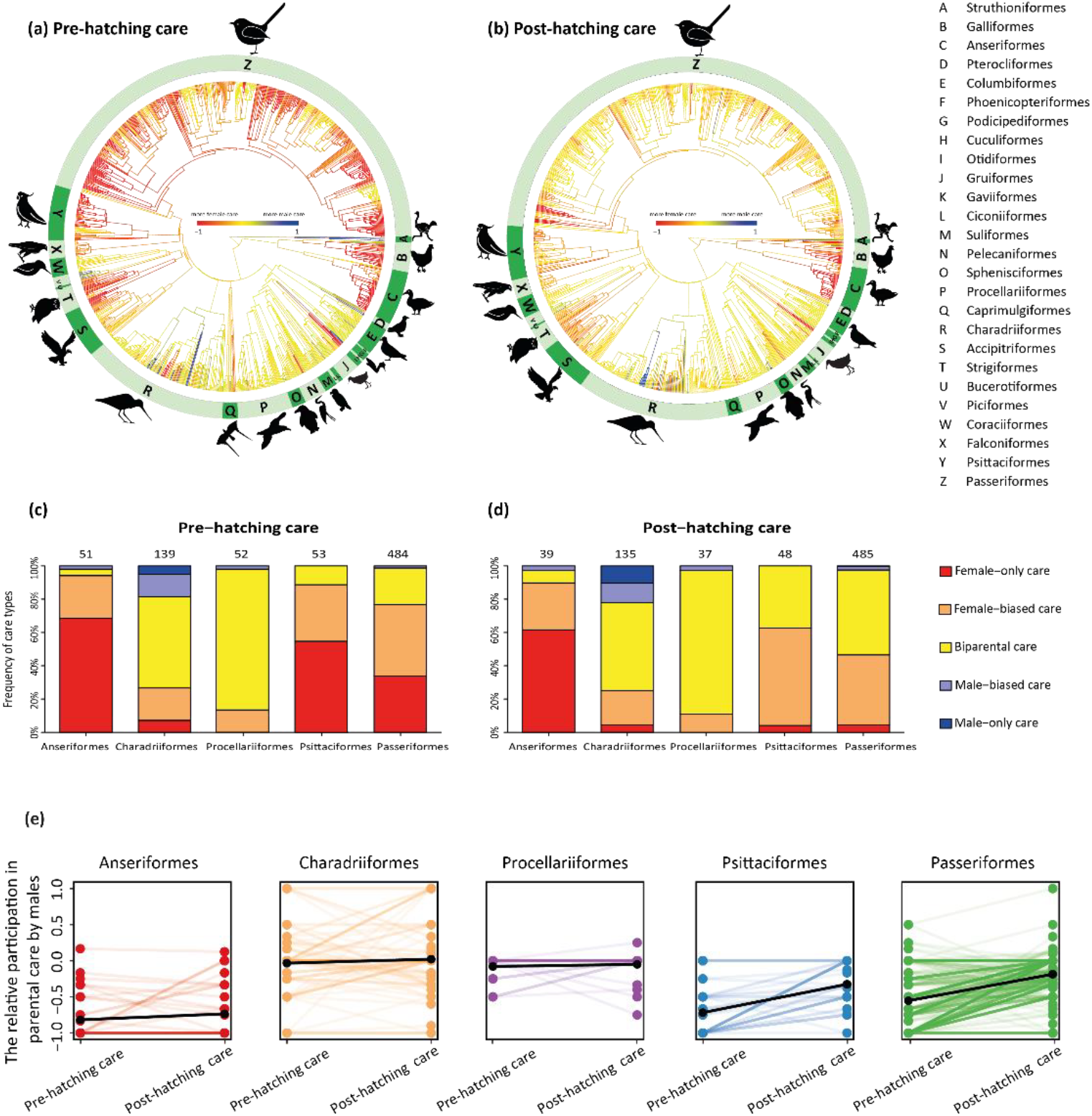
Distribution of parental roles in birds. (**a, b**) Phylogenetic distribution of pre-hatching care and post-hatching care (maximum clade credibility tree of 100 phylogenies using 1065 and 991 bird species, respectively). The figure shows the relative participation in parental care by males for each bird species. (Red = female-only care, yellow = biparental care, blue = male-only care, other colors = sex-biased care). (**c, d**) Frequencies of different parental care patterns in two breeding phases in five major clades. Care patterns are classified into 5 categories: female-only care: 0% male care (red); female-biased care: 1–33% male care (orange); biparental care 34–66% male care (yellow); male-biased care: 67–99% male care (purple); male-only care: 100% male care (blue). Sample size is shown at the top of each column. (**e**) Differences in parental roles between pre-hatching and post-hatching phases in five large clades of birds. Relative male pre-hatching care and post-hatching care are represented by scatterplots, the lines connect two breeding phases for each species and overplotted lines appear as darker lines. The means of pre-hatching care and post-hatching of each avian family are plotted in black.

Female-only care is more common in the pre-hatching phase (26.67% of 1065 species) than in the post-hatching phase (7.87% of 991 species), while biparental care is more prevalent during post-hatching (50.66%) than during pre-hatching (33.43%). In contrast, male-only care is rare in both breeding phases (1.03% and 2.02% in the pre- and post-hatching phase, respectively) (Figure 1). This suggests biparental care is predominant in both the pre-hatching and post-hatching phases, and the parental care offered by the male does not differ from by the female in both breeding phases (PGLS fitted an intercept only (with 100 phylogenies), pre-hatching care: *Slope ± SE* = −0.080 ± 0.188, *P* = 0.649, *n* = 1065 species; post-hatching care: *Slope ± SE* = −0.187 ± 0.134 *P* = 0.170, *n* = 991 species). Furthermore, the relative participation in parental care by males is not different between pre-hatching phase and post-hatching phase (Figure 1 and Table 1). We only found that the relative contribution to post-hatching care by males is marginally larger than in pre-hatching care by males in parrots (Psittaciformes, Figure 1e and Table 1).

**Table 1.** Phylogenetic mean for the parental care differs between pre-hatching phase and post-hatching phase. The difference between the relative participation in pre- and post-hatching care by males is compared in all species and five large clades (Anseriformes, Charadriiformes, Procellariiformes, Psittaciformes and Passeriformes,). Estimates are phylogenetic mean difference with standard error (*Mean difference ± SE*), the corresponding *t* and *p*-values of 100 phylogenetic paired t-test repeated with different phylogenies. log-likelihood of the fitted model *log(L),* phylogenetic signal λ and the number of species *n* are also given for each model.

| Phylogenetic paired t-test | *Mean difference ± SE* | *t* | *p* | *Log(L)* | $\lambda$ | *n* |
| --- | --- | --- | --- | --- | --- | --- |
| **All species** | 0.052 ± 0.184 | 0.340 | 0.735 | -401.4 | 0.626 | 955 |
| **Anseriformes** | -0.080 ± 0.063 | -1.405 | 0.181 | -14.08 | 0.019 | 38 |
| **Charadriiformes** | -0.024 ± 0.111 | -0.221 | 0.826 | -77.62 | 0.334 | 130 |
| **Procellariiformes** | -0.084 ± 0.117 | -0.798 | 0.437 | 0.198 | 0.502 | 35 |
| **Psittaciformes** | **-0.299 ± 0.170** | **-1.921** | **0.079** | **-19.16** | **0.520** | **48** |
| **Passeriformes** | -0.257 ± 0.187 | -1.684 | 0.102 | -154.5 | 0.667 | 459 |

### Diet

Food type is not associated with parental roles: neither parental diet nor nestling diet is associated with sex differences in parental roles (Table 2). In the pre-hatching phase, females provide more care than males no matter what types of food they forage for. While the approximate identical care level is provided by the male and the female in both herbivorous and carnivorous bird species during the post-hatching phase (Figure 2a,b). The lack of relationship between food type and sex-specific parental roles is consistent between the bivariate (Table 2a) and multi-predictor models in which the effects of all potential variables are included (Table 2b,c).

**Table 2.**
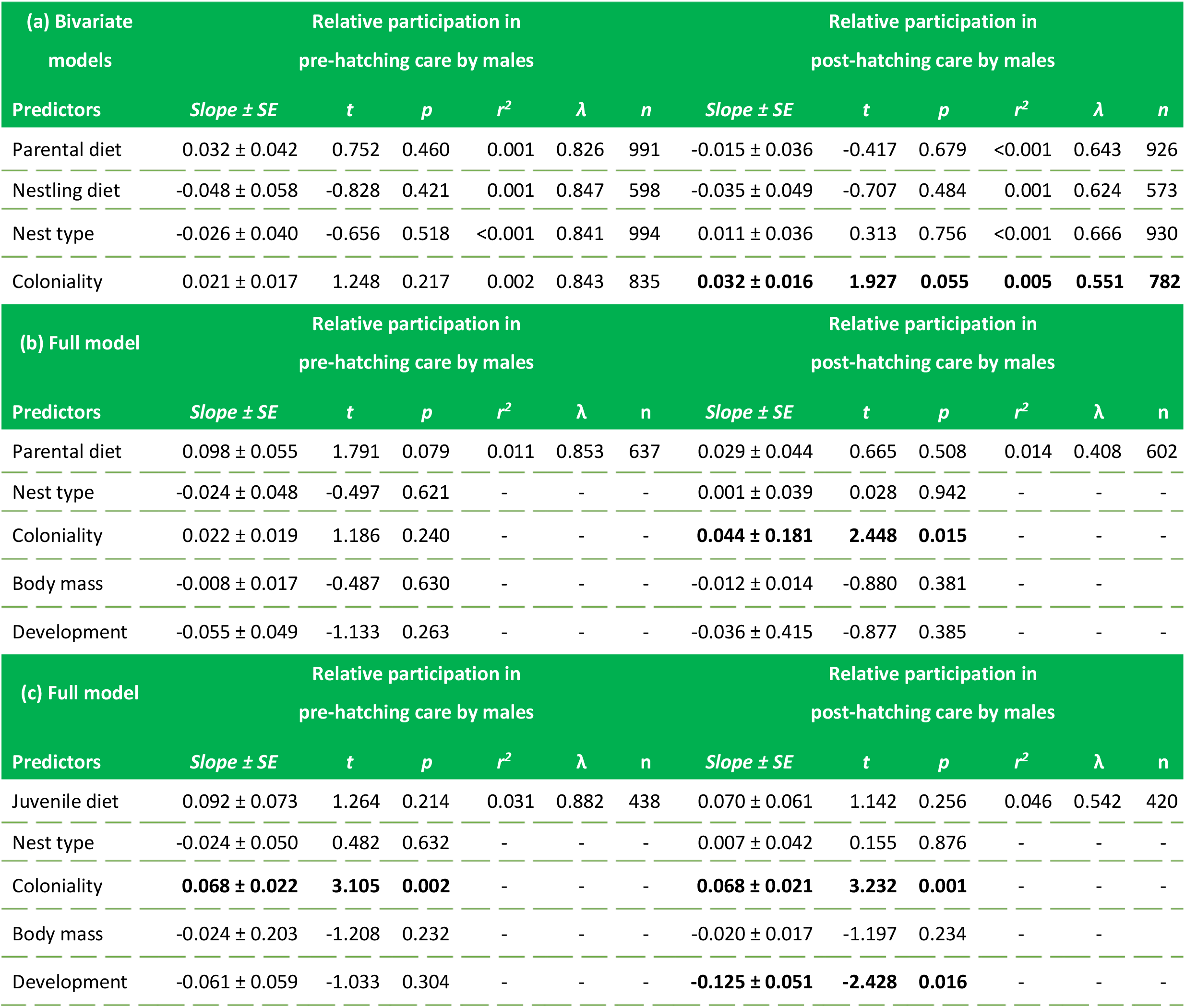
Pre- and post-hatching care in relation to ecology and social environment in birds using phylogenetically generalized linear squares models (PGLS). In both bivariate and multi-predictor PGLS models, the relative participation in pre-hatching care and post-hatching care by males are the response variables, respectively. Predictors include parental diet (plant vs. animal food), nesting diet (plant vs. animal food), nest type (open vs. closed), coloniality (solitary, semi-colonial, colonial). Development mode (precocial vs. altricial) and body mass (log-transformed) are included in the full multi-predictor models. Estimates are means of regression coefficients with standard error (*Slope ± SE*), the corresponding *t* and *p-values* of 100 PGLS analyses repeated with different phylogenies, significant results are highlighted in bold. Sample size *n*, R-squared *r^2^* and phylogenetic signal *λ* are also given for each model.

**Figure 2.**
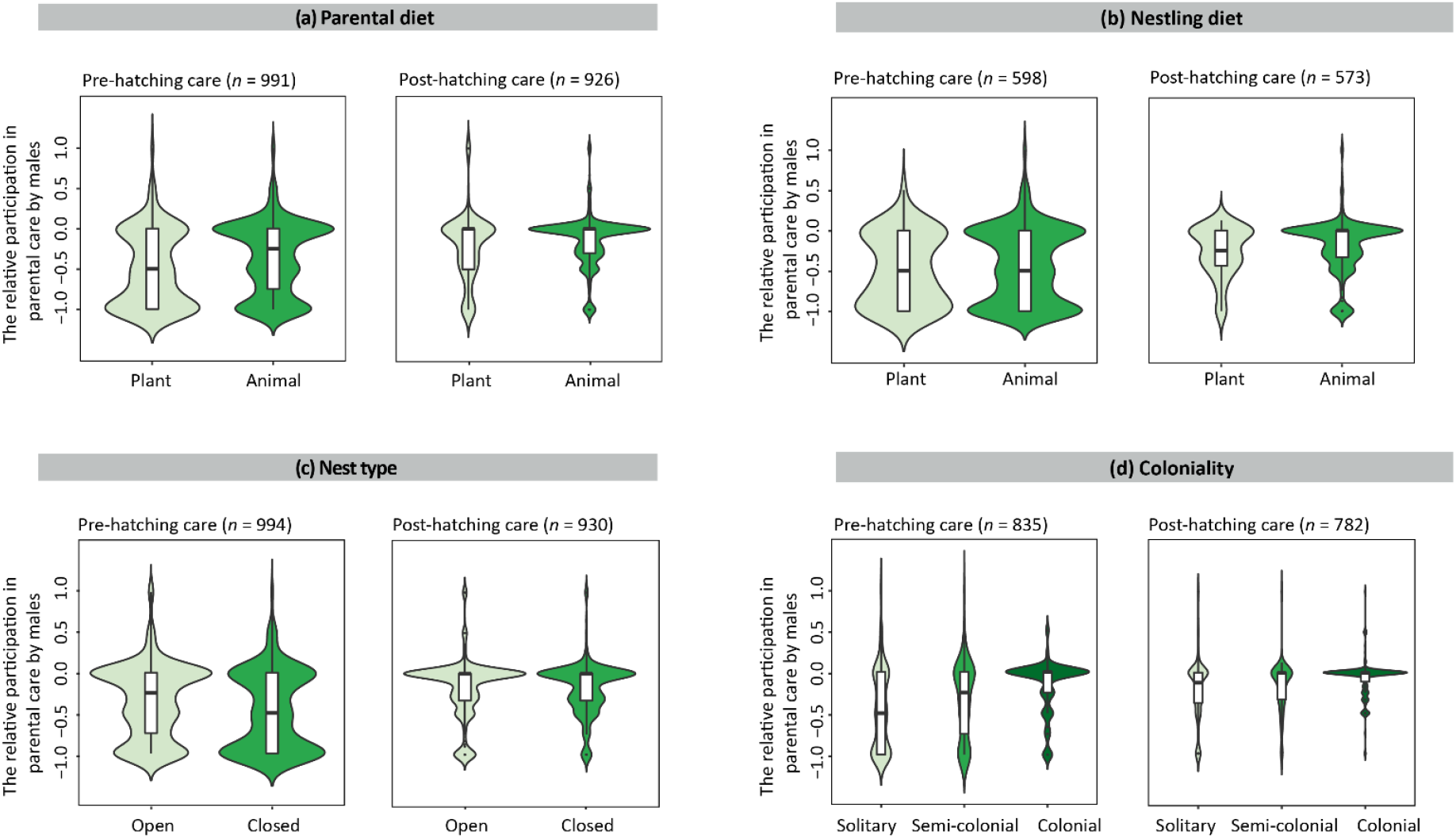
The association between parental roles and ecological and social environment. (a) Parental diet, (b) nestling diet, (c) nest type and (d) coloniality are plotted against relative male pre-hatching care and relative male post-hatching care, respectively. The rectangle of small box plot in each violin plot shows the ends of the first and third quartiles and a vertical line indicates the median value of male care relative to female care. The kernel density plot of each violin plot shows the distribution of parental care and its probability density. The participation in parental care by males was scored on a 5-point scale, −1: 0% male care; −0.5: 1–33% male care; 0: 34–66% male care; 0.5: 67–99% male care; 1: 100% male care. Parental diet and nestling diet were scored on a 2-point scale, 0: plant food, 1: animal food. Nest type was scored on a 2-point scale, 0: open nest, 1: closed nest. Coloniality was scored on a 3-point scale, 0: solitary, 1: semi-colonial, 2: colonial. The number of species n is shown for each plot.

### Nest type

Nest type does not predict parental sex roles, as the relative participation in parental care by males is not significantly different between open and closed nests either in bivariate (Table 2a) or multiple regression analyses (where parental diet and nestling diet are controlled for separately) (Table 2b,c): female-biased pre-hatching care is associated with both open and closed nests, and more egalitarian biparental post-hatching care is found in both nest types (Figure 2c), corresponding to the general care pattern where female-biased care predominates in the pre-hatching phase while biparental care predominates in the post-hatching phase (Figure 1).

### Coloniality

Coloniality is associated with sex differences in parental care. First, coloniality predicts post-hatching care: the relative participation in post-hatching care by males increases as coloniality increases (Table 2, Figure 2d). The relationship is marginally nonsignificant in the bivariate model (Table 2a), and is significant in the full multi-predictor models where parental diet and nestling diet are controlled for respectively (Table 2b,c). However, these fitted PGLS models account for a modest variability of post-hatching care (*r^2^* = 4.6%). Second, relative male pre-hatching care is not related to coloniality (Table 2), although we found a significant trend in the multi-predictor model in which nestling diet instead of parental diet is statistically controlled for (Table 2c).

## DISCUSSION

Our comprehensive phylogenetic comparative analyses confirm that parental roles are highly diverse among avian species, and biparental care is the prevailing care pattern in both pre- and post-hatching phases (Cockburn 2006). Moreover, female-only care is relatively common in the pre-hatching phase, in line with the fact that in approximately 30% of passerine birds only females incubate the eggs (White and Kinney 1974). In addition, the relative participation in parental care by each sex is not remarkably differ between the pre- and post-hatching phases, indicating brood desertion after hatching by either sex happens rarely, but in some precocial bird species such as shorebirds (see Clutton-Brock 1991, Székely and Williams 1995).

The results consistently show that colonial breeding is associated with more equal share in parenting duties than in solitary species, especially in post-hatching care. We think the interaction of two factors could explain the outcome. First, intraspecific competition induces the cooperation between the male and the female parent, since one of the parents has to protect the fragile broods which are completely exposed to the environment, while the other parent competes for food which is used to feed the offspring. Both chick feeding and chick guarding are involved in post-hatching state, therefore, the association between colonial breeding and share in post-hatching care between two sexes is observed in all models. Second, females in a colony synchronously produce the offspring (Gochfeld 1980, Nelson 1980, Coulson 2002), which remarkably reduces the mating opportunities of a deserting male and consequently favors the emergence of biparental care. This outcome demonstrates that equal parental roles are selected under resource and mating constraints in colonial species, whereas female-biased care, mediated by other factors, is favored in solitary species.

Our results demonstrate that nest type does not predict sex-specific parental roles. In the pre-hatching and post-hatching phases, both parents provide care in open nests and closed nests. In the open nests, high predation rates might induce the same response by the male parent and the female parent. Both the male and the female decrease their parental activities around the nests in order to reduce nest visibility, or alternatively, both parents provide more protection against nest predation. In the closed nests with large clutch size, the cooperation between parents is required as large amount of energy and time are needed to raise the offspring. However, we neglected other factors which might be important to explore the relationship between nest habitat and parental roles. For instance, nest sites are essential for breeding success in that good nest sites might promote the development of the offspring (e.g., open nests which are built on the water or in the trees can help to reduce the probability of being predated (Martin 1993; Picman et al. 1993; Colombelli-Négrel and Kleindorfer 2009; Latif et al. 2012)). Further studies investigating the effect of nest microhabitats on sex differences in parental roles will be valuable.

Our results also illustrate that food type cannot explain the considerable variation in parental sex roles: no matter plant or animal food the bird parents forage, the female and the male take care of the offspring together. This suggests both plant-eating and animal-eating offspring might have high demand for food: herbivores request large amount of food while carnivores require high quality but secluded food, inducing biparental care in most of species. Nonetheless, our study did not directly quantify the food availability which is crucial for breeding activities (Martin 1987; Low et al. 2012), since just few empirical data are available. To dig into the impact of diet on parental roles, more empirical studies on food availability are needed in the future. Besides, our study did not take into account the situation where males feed the females during incubation in some avian species (Martin and Ghalambor 1999, Matysioková et al. 2011). Male feeding should be considered as a type of parental care as well, since it increases the survival probability of females and also the ability of females to provide continuous incubation of the eggs, which consequently increases the survival probability of the offspring. The food items which are fed to females might also play an important role in sex role divergence.

In summary, our study provides the most comprehensive analyses of the effect of ecology and social environment on sex differences in parental roles using birds as model organisms. We show that ecological factors such as diet and nest type are not predictors of parental roles, although these factors have strong impacts on some life-history traits. Besides, we found that social environment as coloniality could predict parental care strategies. Further studies including empirical and comparative analyses are needed to explore the relationship between food availability, nest habitats, breeding density and parental sex roles.

## Authors’ contributions

All authors conceived the study. X.L. and A.L. collected the data, X.L. conducted the data analyses with inputs from A.L. All authors interpreted the results. X.L. wrote the manuscript and others contributed important edits.

## ACKNOWLEDGMENTS

We appreciate that Z. Végvári helped with statistical analysis, and we would like to thank the Center for Information Technology of the University of Groningen for their support and for providing access to the Peregrine high performance computing cluster.

## DATA ACCEESBILITY

All relevant data within this paper and its electronic Supplementary material are available once the manuscript is accepted.

