## Supplementary information for "How do ecological and social environments reflect parental roles in birds? A comparative analysis"

Supplementary Figure S1. Scatterplots of different phases of breeding activity in relation to pre- or post-hatching care.

Supplementary Table S1. Pre- and post-hatching care in relation to different types of parental care.

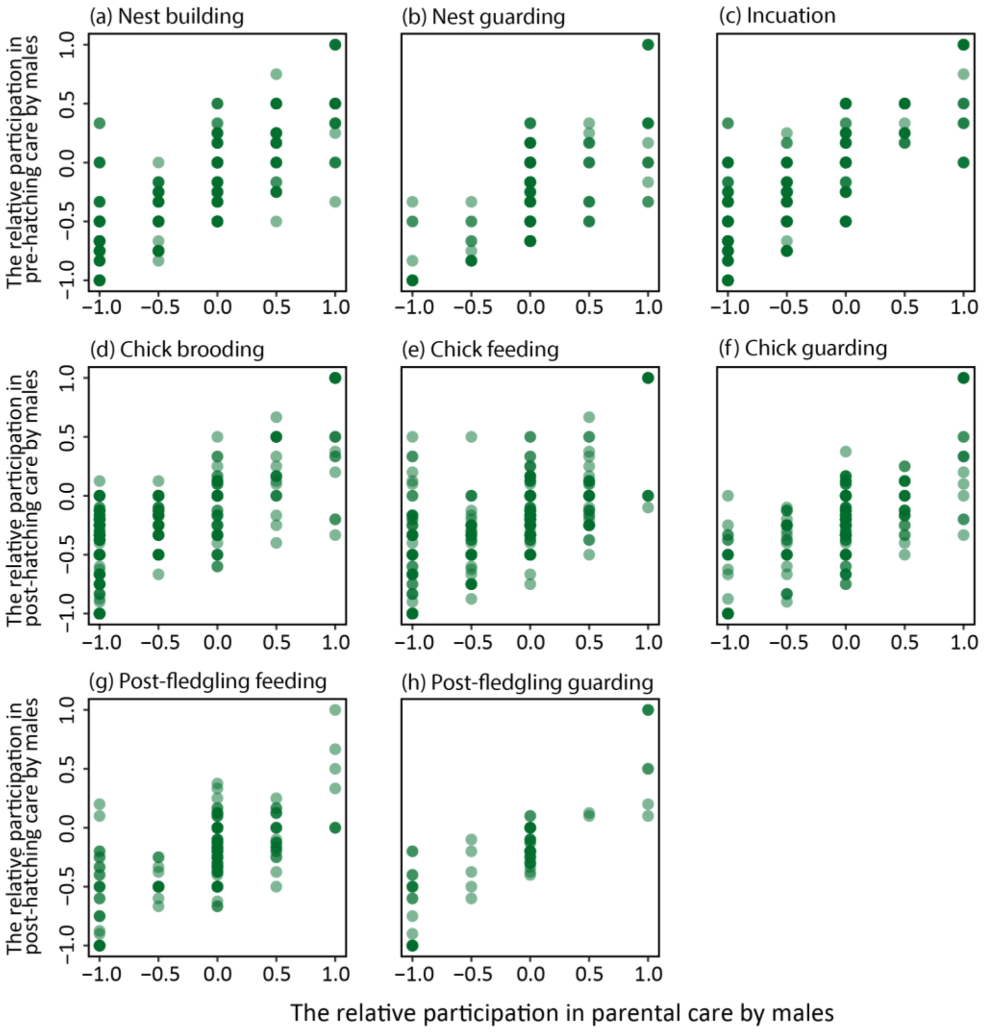


**Figure S1. Scatterplots of different phases of breeding cycle in relation to pre- or post-hatching care.** The relative participation in pre-hatching care by males is associated with the relative contribution to (a) nest building (*n* = 802 species), (b) nest guarding (*n* = 196 species) and (c) incubation by males (*n* = 1017 species). And The relative participation in post-hatching care is correlated with the relative contribution to (d) chick brooding (*n* = 742 species), (e) chick feeding (*n* = 899 species), (f) chick guarding (*n* = 360 species), (g) post-fledgling feeding (*n* = 435 species) and (h) post-fledgling guarding (*n* = 79 species) by males. For each type of parental care, it was scored on a 5-point scale: -1: no male care; -0.5: 1–33% male care; 0: 34–66% male care; 0.5: 67–99% male care; 1: 100% male care. All relationships are significant that *p-value* < 0.001.

**Table S1. Pre- and post-hatching care in relation to different types of parental care.** In PGLS bivariate models, the relative participation in pre-hatching care and post-hatching care by males are the response variables. The predictors include the relative participation in nest building, nest guarding, incubation, chick brooding, chick feeding, chick guarding, post-fledgling feeding and post-fledgling guarding by males. Estimates are means of regression coefficients with standard error (*Slope* ± *SE*), the corresponding *t* and *p*-values of 100 PGLS analyses repeated with different phylogenies. R-squared $r^2$, phylogenetic signal λ and sample size *n* are also given for each model.

| Response variables | Explanatory variables | *Slope ± SE* | *t* | *p* | $R^2$ | λ | *n* |
|---|---|---|---|---|---|---|---|
| **The relative participation in pre-hatching care by males** | The relative participation in nest building by males | 0.006 ± <0.001 | 0.380 | <0.001 | 0.644 | 0.755 | 802 |
| | The relative participation in nest guarding by males | 0.005 ± <0.001 | 0.142 | <0.001 | 0.508 | 0.682 | 196 |
| | The relative participation in incubation by males | 0.007 ± <0.001 | 0.374 | <0.001 | 0.580 | 0.594 | 1017 |
| **The relative participation in post-hatching care by males** | The relative participation in chick brooding by males | 0.005 ± 0.0001 | 0.300 | <0.001 | 0.549 | 0.536 | 742 |
| | The relative participation in chick feeding by males | 0.006 ± <0.001 | 0.320 | <0.001 | 0.533 | 0.566 | 899 |
| | The relative participation in chick guarding by males | 0.006 ± <0.001 | 0.234 | <0.001 | 0.604 | 0.579 | 360 |
| | The relative participation in post fledgling feeding by males | 0.005 ± <0.001 | 0.167 | <0.001 | 0.393 | 0.667 | 435 |
| | The relative participation in post fledgling guarding by males | 0.006 ± <0.001 | 0.144 | <0.001 | 0.729 | 0.187 | 79 |
